# Augmenting chlorophyll fluorescence-based analysis of plant photosynthesis with genetically encoded biosensors

**DOI:** 10.64898/2026.11.24.768924

**Authors:** Ana Paula Cislaghi, Pedro Barreto, Jan-Ole Niemeier, Minh Hoang, Annika Brünje, Gustaf E. Degen, Karin Busch, Felix Buchert, Michael Hippler, Alfred Roland Holzwarth, Markus Schwarzländer

**Affiliations:** Institute of Plant Biology and Biotechnology, University of Münster, Schlossplatz 7/8, 48143 Münster, Germany; Institute of Integrative Cell Biology and Physiology, University of Münster, Schlossplatz 5, 48149 Münster, Germany; Plants, Photosynthesis and Soil, School of Biosciences, University of Sheffield, Firth Court, Western Bank, Sheffield S10 2TN, UK; ChloroSpec B.V., De Boelelaan 1105, 1081 HV Amsterdam, The Netherlands

## Abstract

The status of the light reactions of photosynthesis is tightly linked to subcellular physiology and metabolism. Yet, monitoring how both these levels are coordinated in living plants has remained challenging to explore. Here, we combine genetically encoded biosensors with time-and wavelength-resolved chlorophyll fluorescence spectroscopy (ChloroSpec L1) in attached *Arabidopsis thaliana* leaves. Stromal pHluorin2 reported a reversible, light-dependent alkalinization in a single mesophyll layer of leaf discs imaged by confocal microscopy and over the illuminated area of attached leaves recorded by ChloroSpec, where a distinct overshoot appeared at all irradiances. Measurement of the light-response dynamics of stromal MgATP^2-^, cytosolic pH, and cytosolic NADH/NAD^+^ illustrates the flexibility of the approach to different biosensors and subcellular localizations. Under prolonged fluctuating light, simultaneous recordings linked progressive stromal alkalinization and slow cytosolic NAD redox integration to wavelength-dependent shifts in photosystem II efficiency and non-photochemical quenching. By coupling programmable illumination with multiplexed metabolic and spectroscopic readouts *in planta*, this platform bridges cellular bioenergetics and leaf physiology to dissect photosynthetic regulation under dynamic environments.

## Introduction

Photosynthesis couples light capture to proton translocation across the thylakoid membrane, which alkalinizes the stroma (Heldt et al., 1973) and, through acidification of the thylakoid lumen, drives ATP synthesis, alongside NADP^+^ reduction. The resulting ATP and NADPH fuel CO_2_ assimilation through the Calvin–Benson–Bassham (CBB) cycle. Photosynthetic activity can also influence the redox state of metabolites beyond the chloroplast through the transfer of reducing equivalents to the cytosol (Shameer et al., 2019; Steinbeck et al., 2020; Zheng et al., 2025). Under natural conditions, irradiance changes continuously, and the regulation of these processes on time scales from seconds to hours determines photosynthetic efficiency (Armbruster et al., 2017; Kaiser et al., 2018). Following the dynamics of pH, energy and redox state in living tissue is therefore essential to understand how photosynthesis is regulated.

Genetically encoded biosensors enable the monitoring of such parameters *in vivo* and in defined subcellular compartments, as demonstrated for pH with pHluorin2 (Mahon, 2011; Cislaghi et al., 2026), MgATP^2-^ with ATeam1.03-nD/nA (Imamura et al., 2009; De Col et al., 2017) and NADH/NAD^+^ with Peredox-mCherry (Hung et al., 2011; Steinbeck et al., 2020). In plants, these sensors have mostly been read by confocal laser scanning microscopy (CLSM) of leaf discs, which resolves individual cell layers and, combined with on-stage illumination devices, has revealed light-induced changes in several compartments (Elsässer et al., 2020; Zheng et al., 2025). However, this approach is restricted to excised tissue and does not simultaneously provide photosynthetic parameters, such as the effective quantum yield of photosystem II (PSII), Y(II), and non-photochemical quenching, NPQ, which are calculated from chlorophyll fluorescence and report on photosynthetic performance (Genty et al., 1989; Kramer et al., 2004; Klughammer & Schreiber, 2008; Baker, 2008).

ChloroSpec L1 (ChloroSpec B.V., Amsterdam, The Netherlands) is a spectrometer-based system that records wavelength- and time-resolved fluorescence from attached leaves under programmable illumination, while simultaneously deriving photosynthetic parameters from chlorophyll fluorescence (Nanda et al., 2024). Because the sample can be excited at different wavelengths and the emission spectrum is collected rapidly over a broad range, ChloroSpec can also capture changes of a biosensor together with these parameters in the same leaf. However, it remains unclear how the biosensor kinetics measured with ChloroSpec compare with those obtained by CLSM, and whether recording both types of readout simultaneously can link metabolic changes to photosynthetic performance.

Here, we compare the stromal pH responses of pHluorin2 measured by CLSM and by ChloroSpec, and we use the latter to analyse the biosensor response under different light regimes. We further show that the approach can be extended to MgATP^2-^ and NADH/NAD^+^ dynamics, providing a basis for relating biosensor readouts to photosynthetic performance in intact plants.

## Results

### Confocal microscopy and ChloroSpec both resolve light-induced stromal pH dynamics

To compare the two platforms, we used *Arabidopsis thaliana* plants expressing the ratiometric pH biosensor pHluorin2 in the chloroplast stroma (Cislaghi et al., 2026). *In vitro*, purified pHluorin2 has two excitation maxima, at 387 and 482 nm, and a single emission peak at 510 nm (Mahon et al., 2011; Cislaghi et al., 2026). We excited it close to these maxima, at 405 nm on both platforms and at 488 nm in CLSM or 465 nm in ChloroSpec. With increasing pH, the emission obtained on excitation in the shorter-wavelength band increases, whereas that obtained on excitation in the longer-wavelength band decreases. The ratio between the two channels therefore reports stromal pH, with a higher ratio indicating alkalinization (Fig. 1A), so that the responses are described hereafter in terms of pH.

**Figure 1.**
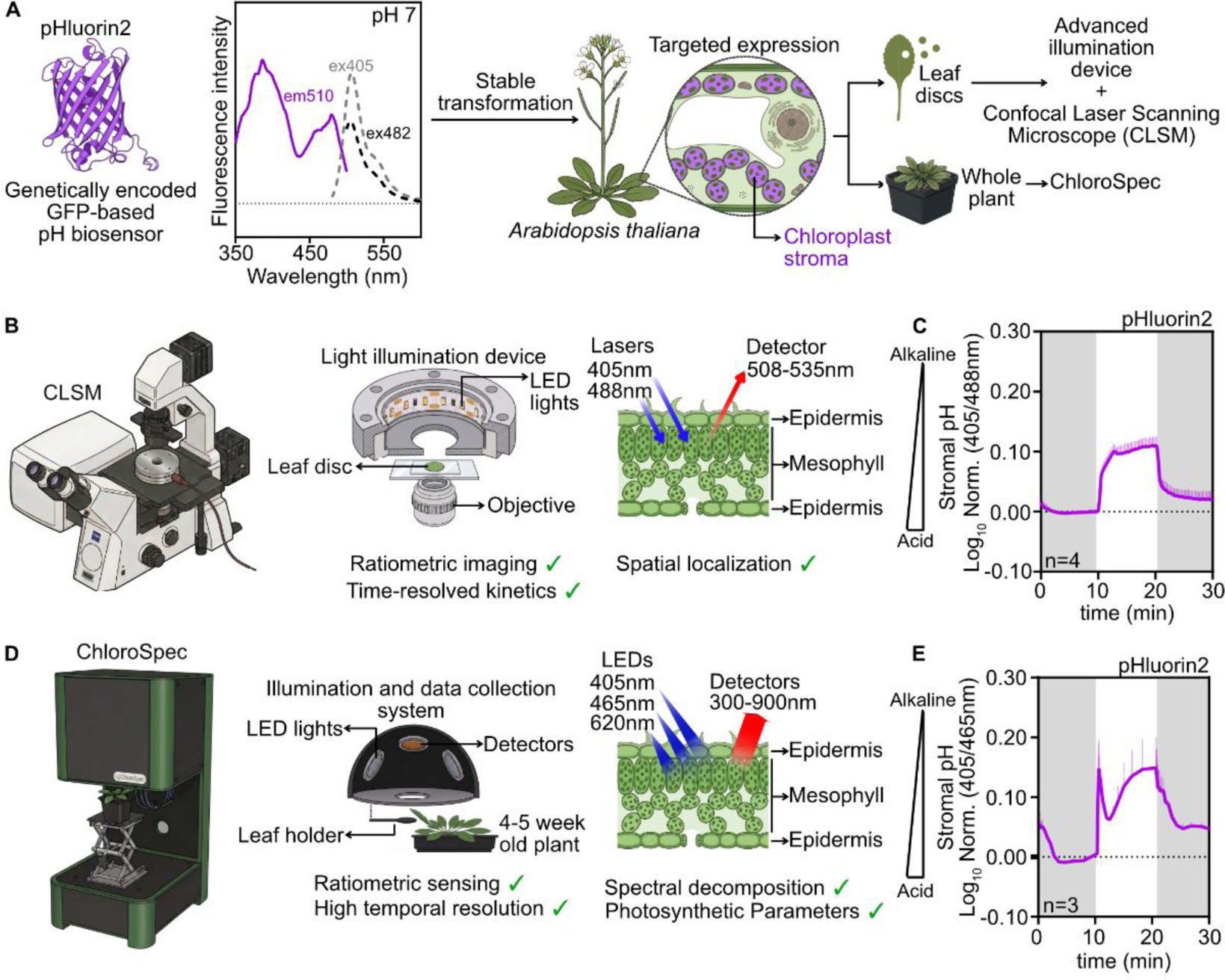
Dual platforms for *in vivo* monitoring of chloroplast stromal pH dynamics in *Arabidopsis thaliana*. **A**, Experimental workflow. pHluorin2, a genetically encoded ratiometric pH biosensor based on GFP, was stably expressed in the chloroplast stroma. Spectra of pHluorin2 at pH 7, adapted from Cislaghi et al., 2026. The solid line is the excitation spectrum collected at an emission of 510 nm, and the dashed lines are the emission spectra obtained on excitation at 405 nm and at 482 nm. Leaf discs were analysed by CLSM with a programmable LED illumination device, and attached leaves of intact plants were analysed with ChloroSpec. **B,** CLSM platform. Leaf discs were placed on a custom LED illumination device mounted on the microscope stage. pHluorin2 was excited sequentially with 405 nm and 488 nm lasers, and emission was collected between 508 and 535 nm. **C,** Stromal pH kinetics measured by CLSM in leaf discs during 10 min of dark, 10 min of light at 100 µE and 10 min of dark, with grey shading indicating dark and white indicating light. The y-axis shows the normalized log_10_ of the 405/488 nm fluorescence ratio. Data are mean ± s.d. of *n* = 4 leaf discs, each from a different plant. **D,** ChloroSpec platform. An attached leaf of an intact 4–5-week-old plant was held in the leaf holder of an integrated illumination and detection system with three LEDs, here at 405, 465 and 620 nm and a detector covering 300–900 nm. This allows ratiometric biosensor measurements with high temporal resolution together with wavelength-resolved recording of photosynthetic parameters *in planta*. pHluorin2 was excited at 405 and 465 nm, and the ratio was calculated from emission at 511 nm. **E,** Stromal pH kinetics measured with ChloroSpec in attached leaves under the same light regime as in C. The y-axis shows the normalized log_10_ of the 405/465 nm fluorescence ratio. Data are mean ± s.d. of *n* = 3 leaves, each from a different plant.

The CLSM is equipped with an on-stage illumination device, a ring of LEDs that exposes the sample to defined light intensities and durations (Zheng et al., 2025). This configuration allows individual cell layers of a leaf disc to be imaged during illumination, with pHluorin2 excited by 405 nm and 488 nm lasers and its emission collected between 508 and 535 nm. To follow pH changes during the activation of photosynthesis, we imaged the mesophyll layer, which contains the chloroplasts that carry out most of the photosynthetic activity of the leaf (Terashima et al., 2011) (Fig. 1B). ChloroSpec, in contrast, excites the sample with three LEDs. One of them (620 nm) excites chlorophyll fluorescence, from which photosynthetic parameters such as those obtained by pulse-amplitude-modulation (PAM) fluorometry are calculated, whereas the other two can be adapted to the biosensor in use (here 405 and 465 nm for pHluorin2). Emission is detected simultaneously across a broad wavelength range of 300–900 nm. In addition, a leaf of an intact plant can be held between two clips close to the LEDs and the detectors, so that the signal is averaged over the illuminated leaf area, integrating multiple cell layers, rather than within a single cell layer as in CLSM, which also makes the readout comparable to measurements with DUAL-PAM and DUAL-KLAS-NIR instruments.

Under the same dark–light–dark regime of 10 min each, with a light intensity of 100 µE (µmol photons m^-2^ s^-1^), both platforms detected a reversible, light-dependent increase of stromal pH, consistent with the light-driven proton flux into the thylakoid lumen (Heldt et al., 1973; Trinh & Masuda, 2022). In leaf discs, pH increased rapidly on illumination and reached a plateau that was maintained until the light was switched off, when it decreased rapidly to values close to those before illumination, as described for low and moderate intensities (Zheng et al., 2025; Cislaghi et al., 2026) (Fig. 1C). In attached leaves, by contrast, the response was biphasic: pH rose rapidly to an initial maximum (hereafter overshoot), decreased partially, indicating partial re-acidification, and rose again to a plateau that persisted until the light was switched off (Fig. 1D,E). This profile could reflect a transient excess of proton influx over efflux before the proton-efflux pathways are fully engaged. In CLSM, a biphasic pattern appeared only at the higher intensities of 200 and 850 µE (Zheng et al., 2025; Cislaghi et al., 2026) (Fig. S1).

Both instruments resolved a reversible, light-dependent alkalinization of the stroma, but the shape of the response at 100 µE and the dependence of its amplitude on light intensity differed (Figs. 1, 3B and S1), so that the two platforms provide complementary rather than equivalent views of the same process.

### ChloroSpec resolves light-induced pH, MgATP^2-^ and NADH/NAD^+^ dynamics in different compartments

Photosynthesis alters not only stromal pH but also the energy and redox balance across multiple subcellular compartments (Kramer & Evans, 2011; Scheibe, 2004). To test whether ChloroSpec can monitor biosensors with distinct spectral properties in different compartments, we recorded plants expressing pHluorin2 in the cytosol, ATeam1.03-nD/nA (MgATP^2-^; De Col et al., 2017) in the stroma and Peredox-mCherry (NADH/NAD^+^; Steinbeck et al., 2020) in the cytosol during dark–light–dark regimes at 100 µE (Fig. 2). Each sensor reported a distinct light response.

**Figure 2.**
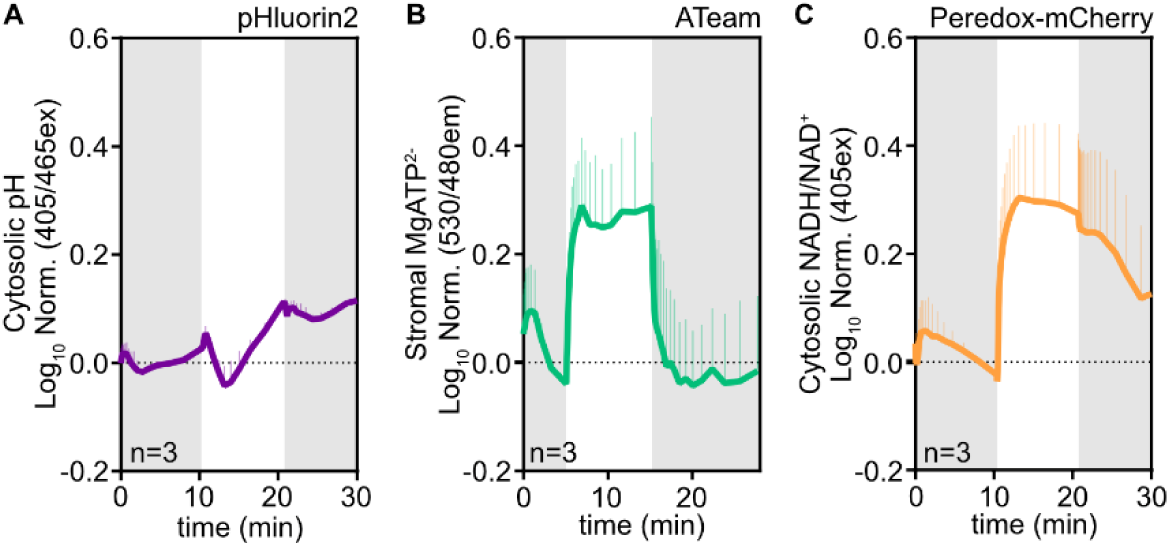
Monitoring of subcellular pH, MgATP^2-^ and NADH/NAD^+^ dynamics during illumination with ChloroSpec. **A**, Cytosolic pH measured with pHluorin2 (normalized log_10_ of the 405/465 nm ratio) in attached leaves during 10 min of dark, 10 min of light at 100 µE and 10 min of dark (grey shading, dark; white, light). **B,** Stromal MgATP^2-^ measured with ATeam1.03-nD/nA (normalized log_10_ of the 530/480 nm emission ratio, 440 nm excitation) under 100 µE, with 5 min of darkness, 10 min of light and about 13 min of darkness. **C,** Cytosolic NADH/NAD^+^ measured with Peredox-mCherry (normalized log_10_ of tSapphire fluorescence, 405 nm excitation, 520 nm emission) under the light regime in A. Data are mean ± s.d. of *n* = 3 leaves, each from a different plant.

Cytosolic pH showed a brief alkalinization within the first minute of light, followed by acidification below the pre-illumination level after about 3 min and then a gradual alkalinization to a level that persisted after light-off (Fig. 2A). Cytosolic alkalinization may reflect the removal of protons by thylakoid energization and by transfer into the vacuole (Felle & Bertl, 1986; Yin et al., 1990a, 1990b), whereas the later phases may depend on carbon metabolism (Yin et al., 1990a). A higher cytosolic pH after illumination than before has also been reported with cpYFP (Elsässer et al., 2020).

Because the same proton gradient drives ATP synthesis, stromal MgATP^2-^ also responded to illumination. It rose rapidly and reached a maximum after about 2 min of light, decreased slightly and then remained well above the dark level until the light was switched off, when it fell rapidly to about the pre-illumination value (Fig. 2B). The initial increase most likely reflects light-driven ATP synthesis, since ATP import into mature chloroplasts is negligible (Voon et al., 2018), and the slight decrease could reflect the activation of ATP-consuming processes such as the CBB cycle (Voon et al., 2018).

Photosynthesis also generates reducing power alongside ATP. The cytosolic NADH/NAD^+^ signal increased steeply within about 3 min of illumination to a plateau and decreased slowly in the dark without returning to the initial level (Fig. 2C). This is consistent with the export of reducing equivalents from the chloroplast (Steinbeck et al., 2020), probably through the malate valve and the triose phosphate–3-phosphoglycerate shuttle (Shameer et al., 2019). A brief oxidation after the initial increase has been resolved by CLSM in leaf discs, whereas we did not detect it with ChloroSpec (Zheng et al., 2025).

Thus, the three sensors reported distinct kinetics. Stromal MgATP^2-^ rose within about 2 min of illumination and decreased rapidly at light-off, cytosolic pH changed in several phases over minutes and remained elevated in darkness, and cytosolic NADH/NAD^+^ increased within about 3 min and relaxed slowly in darkness. These different kinetics show that ChloroSpec resolves compartment- and metabolite-specific responses with biosensors of different spectral properties.

### ChloroSpec resolves stromal pH responses to different irradiances within a single recording

A distinctive feature of ChloroSpec is programmable sequential illumination within a single recording, which allows the dependence of the biosensor response on light intensity to be examined in one experiment. On the confocal platform, illumination is triggered by the microscope software and the light intensity is set with a dimmer (Zheng et al., 2025), whereas ChloroSpec allows the illumination protocol to be defined in advance, so that the intensity and duration of every phase are applied automatically (Nanda et al., 2024). We therefore exposed attached leaves of independent plants expressing stromal pHluorin2 to alternating 5-min dark and light phases of increasing intensity, from 25 to 800 µE, while recording the full emission spectrum throughout (Fig. 3A, Fig. S2A–C).

**Figure 3.**
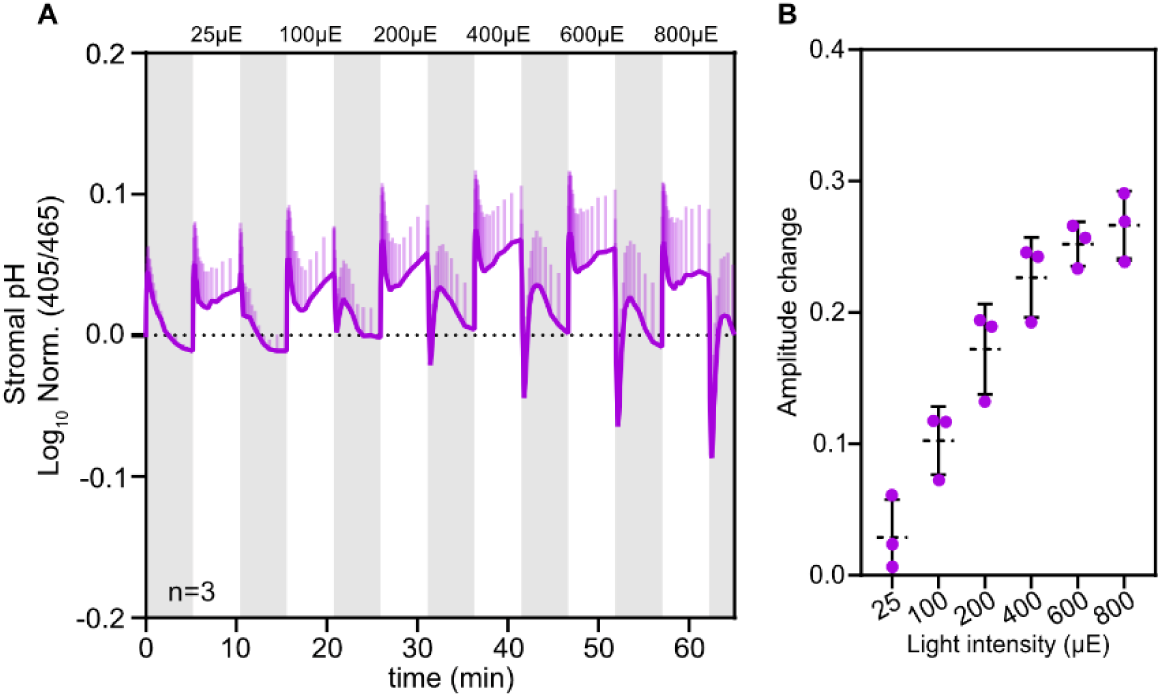
Light dose-dependent *in planta* chloroplast stromal pH dynamics. **A**, Stromal pH measured with pHluorin2 as the normalized log_10_ of the 405/465 nm ratio in attached leaves exposed to six 5-min light periods of increasing intensity (25, 100, 200, 400, 600 and 800 µE; white bars), alternating with 5-min dark phases (grey bars). Data are mean ± s.d. of *n* = 3 leaves, each from a different plant. **B,** Amplitude of the stromal pH response, defined as the difference between the maximum pH after light onset and the minimum after light-off, as a function of light intensity. Points show individual leaves, each from a different plant (*n* = 3), and dashed lines and error bars show mean ± s.d.

In the leaves analysed, each light phase elicited the same response on illumination: the stromal pH increased rapidly to an overshoot, decreased partially and then remained at, or rose slowly towards, a plateau below the overshoot level until the light was switched off. The overshoot was present at all light intensities tested, including low intensities such as 25 µE, which suggests that it is an intrinsic feature of the stromal pH response after darkness and not a consequence of a particular light dose. At light-off, pH decreased sharply and, from 200 µE upwards, fell below the pre-illumination level (undershoot), which became more pronounced at higher intensities, and relaxed towards the pre-illumination level within the following 5 min of darkness (Fig. 3A). The amplitude of the response, defined as the difference between the maximum pH reached after light onset and the minimum reached after light-off, increased with light intensity across the range tested, although the increments became smaller at 600 and 800 µE (Fig. 3B).

### Fluctuating illumination with ChloroSpec reveals wavelength-dependent photosynthetic parameters alongside stromal pH dynamics

Light in nature is rarely constant, because clouds and canopy movement continuously change irradiance, and the slow adjustment of photoprotection and carbon fixation to these changes limits photosynthetic efficiency (Kaiser et al., 2018; Kromdijk et al., 2016). Dynamic regulation of the thylakoid proton motive force is central to this response (Armbruster et al., 2017). ChloroSpec allows the actinic light to be programmed freely (Nanda et al., 2024), which made it possible to expose samples to fluctuating light for extended periods of time. We therefore exposed leaves of plants expressing stromal pHluorin2 to a fluctuating light regime of about 4 h. The regime started with 15 min of constant illumination at 150 µE, followed by sinusoidal modulation between 50 and 250 µE. The period of the oscillations lengthened progressively from about 2 min to 11 min towards the middle of the series and shortened again to about 3 min before the light was turned off (Fig. 4A).

**Figure 4.**
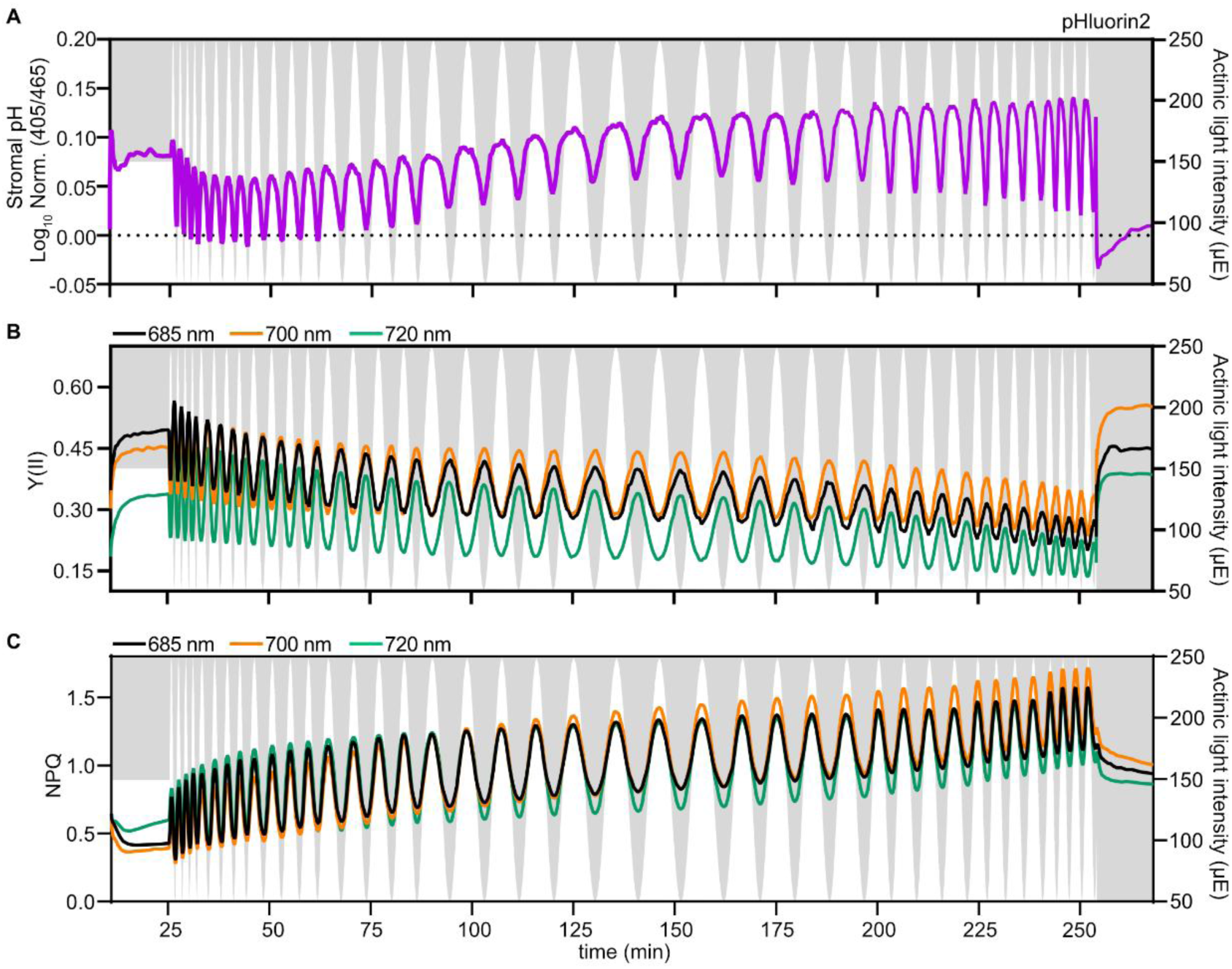
Stromal pH and wavelength-resolved photosynthetic parameters during fluctuating illumination. **A**, Stromal pH measured with pHluorin2 as the normalized log_10_ of the 405/465 nm ratio in attached leaves during a fluctuating light regime of about 4 h. The regime started with 15 min of constant light at 150 µE, followed by sinusoidal modulation between 50 and 250 µE with a period of about 2 to 11 min, and ended in darkness. Time is counted from the start of the recording; the first 10 min, in darkness, are not shown. The actinic light intensity is read on the right y-axes, with white regions indicating illumination and grey regions the absence of light. **B,** Effective quantum yield of photosystem II, Y(II), calculated from fluorescence at 685 nm, shown in black, at 700 nm, in orange, and at 720 nm, in green, in the same leaf as in A. **C**, NPQ calculated from the same fluorescence signals as in B. Traces show one representative leaf of three independent experiments, each from a different plant.

At the onset of illumination, the stromal pH rose rapidly to the overshoot and then declined partially, as observed in the dark–light–dark cycles (Figs. 1 and 3). During the subsequent fluctuations, the stromal pH increased and decreased in phase with irradiance, and the mean pH of the oscillations increased progressively over the series (Fig. 4A). After the last light phase, pH dropped below its initial value and then recovered slowly, consistent with the stromal acidification that follows illumination (Zheng et al., 2025), whereas NPQ decreased only partly and remained above the values at the start of the recording (Fig. 4C).

Notably, in contrast to the overshoot observed at the first illumination after darkness (Figs. 1 and 3), no overshoot appeared after each increase in irradiance during the fluctuations (Fig. 4A), where pH followed a smooth sigmoidal course. A possible explanation is the activation state of the proton-releasing pathways. During the fluctuations, irradiance never decreased below 50 µE, so that the ATP synthase (Kohzuma et al., 2017) and the CBB cycle (Kaiser et al., 2018) probably remained partly active, which could allow stromal pH to follow the photosynthesis-driven alkalinization without an overshoot.

Photosynthetic parameters are conventionally derived from fluorescence detected at wavelengths above about 700 nm, as in standard PAM fluorometers with far-red detection (Pfündel, 2021), yet the values obtained depend on the emission wavelength at which fluorescence is detected (Genty et al., 1990; Stroch et al., 2005; Pfündel, 2021). Because ChloroSpec records the whole emission spectrum in the same measurement, we compared the photosynthetic parameters obtained at detection wavelengths of 685, 700 and 720 nm (Fig. 4B,C). From these signals we calculated Y(II), NPQ, and the yields of regulated, Y(NPQ), and non-regulated, Y(NO), energy dissipation (Genty et al., 1989; Kramer et al., 2004; Klughammer & Schreiber, 2008). Y(II) varied inversely with irradiance and declined over the series, whereas NPQ varied in phase with irradiance and increased (Fig. 4B,C). Y(NPQ) increased while Y(NO) remained nearly constant (Fig. S3A), indicating that the loss of Y(II) was compensated mainly by regulated dissipation. The progressive increase of stromal pH alongside NPQ is consistent with a larger proton gradient (Müller et al., 2001; Avenson et al., 2005).

Detection wavelength affected all parameters, and the differences between wavelengths changed with time (Fig. 4B,C, Fig. S3A). In the first constant-light phase, at about 10 to 25 min, Y(II) decreased with wavelength in the order 685 > 700 > 720 nm, whereas NPQ and Y(NPQ) were highest at 720 nm. In the later part of the recording, Y(II) and NPQ were highest at 700 nm and lowest at 720 nm, whereas Y(NPQ) and Y(NO) were highest at 720 nm and lowest at 700 nm. Some difference between detection wavelengths is expected, because emission above about 700 nm contains a larger share of non-variable photosystem I (PSI) fluorescence (Genty et al., 1990; Franck et al., 2002, 2005; Pfündel, 2021), whereas emission below 700 nm is more strongly reabsorbed (Pfündel, 2021), and detection mainly of PSII emission (660–710 nm) yields higher NPQ and PSII efficiencies than standard far-red detection (Stroch et al., 2005). The lower NPQ at 720 nm than at shorter wavelengths in the later part of the recording agrees with earlier ChloroSpec measurements (Nanda et al., 2024; Cainzos et al., 2026), whereas the order was reversed in the first constant-light phase. Wavelength-integrated detection merges these differences into a single value. Such differences are also relevant for linking leaf-level PAM measurements to solar-induced fluorescence, which is retrieved in narrow bands near 687 and 760 nm (Magney et al., 2017).

In addition, we measured cytosolic NADH/NAD^+^ with Peredox-mCherry in separate plants under the same modulated light regime (Fig. S3B). The NADH/NAD^+^ signal reached a maximum at about 45 to 50 min, that is about 20 to 25 min after the start of the modulation, decreased partially over the following 1.5 h, increased again towards the end of the series and decreased after the last light phase. Unlike stromal pH, it did not follow individual fluctuations, which is consistent with the integration of reductant export from the chloroplast (Steinbeck et al., 2020).

Because irradiance oscillated within the same range throughout the regime, the progressive increase of stromal pH and NPQ and the decline of Y(II) may reflect a dependence of the state of the chloroplast on the preceding light history and not only on the instantaneous irradiance.

## Discussion

Our results show that leaf discs measured by CLSM and attached leaves measured with ChloroSpec both report a reversible, light-dependent alkalinization of the stroma. Each technique, however, provides complementary information. CLSM allows biosensor changes to be followed with spatial resolution, within a given cell layer of the leaf and even in individual chloroplasts, whereas ChloroSpec averages over the illuminated leaf area. ChloroSpec, in turn, allows the biosensor response to be examined under different illumination regimes and, in parallel, the photosynthetic parameters to be determined, which relates the measured metabolite to the photosynthetic performance of the plant. The two approaches are therefore complementary, and both are useful to link cellular metabolism to plant physiology.

When comparing the results from the two instruments, it is important to consider their differences. For stromal pH measured with pHluorin2, the curves obtained with each platform were not identical: ChloroSpec detected an overshoot already at 25 µE (Fig. 3A), whereas in the confocal microscope it appeared in the mesophyll only at 200 and 850 µE (Fig. S1), and the rise in pH appeared smaller at 850 than at 200 µE, in contrast to ChloroSpec, where the amplitude increased with intensity (Fig. 3B). Light is attenuated across the leaf, so that cell layers are exposed to different irradiances (Karabourniotis et al., 2021), and chloroplasts near the illuminated surface saturate at lower irradiances than those deeper in the leaf (Terashima et al., 2011). A signal that includes the more strongly illuminated layers could therefore show at 100 µE (Fig. 1C,E) a response that the mesophyll layer imaged by CLSM reaches only at higher intensity. In addition, the light sources differ between the two setups, and a nominal 100 µE in the confocal microscope may not correspond exactly to 100 µE in ChloroSpec, because the intensities were not measured with the same light meter. The physiological state of attached leaves and excised discs may also differ, for instance through wounding and the limited gas exchange of a disc between coverslips.

The kinetics measured in other compartments are consistent with earlier work. The fast increases of stromal MgATP^2-^ and cytosolic NADH/NAD^+^ agree with the reported behaviour of these sensors and with the proposed export of reducing equivalents from the chloroplast (Lim et al., 2020; Steinbeck et al., 2020; Zheng et al., 2025). A pronounced reduction of the cytosolic NAD pool in illuminated leaf discs has also been reported with Peredox in the NAPstar study (Scherschel et al., 2024). Whether the brief oxidation resolved by CLSM (Zheng et al., 2025) reflects sensor-specific behaviour or metabolism, and why it was not detected here, remains to be resolved. That sensors with different spectral properties could be read in the same instrument supports the use of ChloroSpec for further sensors. The biosensor module has already been used with redox sensors (Hipsch et al., 2025), and our work extends it to pH, ATP and NADH/NAD^+^.

Programmable illumination allowed us to apply light regimes closer to those found in nature, although still simplified relative to natural sunlight. When leaves were exposed to different stable irradiances, the overshoot recurred at every intensity, which may point to regulation of the proton circuit at the dark-to-light transition rather than a simple dose response (Armbruster et al., 2017). This profile resembles the kinetics of the P515 signal during the induction of photosynthesis (Schreiber & Klughammer, 2008), and a transient early maximum of the proton motive force can also be seen in wild-type Arabidopsis after a step to high light (Degen et al., 2023). The overshoot may therefore be related to the Kautsky effect and reflect the transient interplay of ATP synthase activation, carbon fixation and NPQ during induction. The smaller increments of the amplitude at high light could be in line with mechanisms that limit the proton gradient or dissipate excess energy, such as photosynthetic control and NPQ (Müller et al., 2001; Rott et al., 2011).

The stepwise and the fluctuating regimes differ in three respects that matter for interpreting the pH response: the starting level, that is whether the leaf returns to darkness, the rate of change and the step size. In the stepwise series, every increase in irradiance was a step that began from a dark-adapted state and produced an overshoot, a lower plateau and, at light-off, a decline to or below the pre-illumination level, so that each cycle was a full dark–light transition whose amplitude scaled with intensity. In the fluctuating regime, only the first illumination began from darkness and showed the overshoot, whereas the later increases in irradiance, which started from 50 µE and never from darkness, followed slow sinusoidal ramps with smaller increments and produced a smooth rise without an overshoot. The pH therefore depended on the type of transition and not only on the irradiance reached. The difference in the starting level fits current views on the regulation of the proton circuit: in darkness, the chloroplast ATP synthase is largely inactive because its γ-subunit is oxidized, which limits proton efflux until it is reduced by thioredoxin in the light (Kohzuma et al., 2017), and the activation of carbon fixation takes minutes (Kaiser et al., 2018). Under fluctuating light that does not experience darkness, both are expected to remain partly active, so that proton efflux can follow the rise in proton influx. The thylakoid K^+^/H^+^ antiporter KEA3 has been shown to accelerate the relaxation of the proton gradient after transitions to lower light (Armbruster et al., 2014), a role that is more relevant in fluctuating than in constant light and that could contribute to the in-phase decreases of pH during the oscillations.

A second difference concerns the level of the signal over time. In the stepwise series pH relaxed after light-off towards its pre-illumination level within the 5 min of each dark phase, so that each light phase started from a similar state and the level reached depended on intensity. In the fluctuating regime the mean level of the oscillations increased progressively although irradiance oscillated within the same range of 50–250 µE, and NPQ rose and Y(II) fell over the same period. This pattern could reflect the accumulation of slowly relaxing NPQ components and is similar to the slow, cumulative build-up of photoprotection described under fluctuating light, in which NPQ and its slow components change on time scales of minutes to hours (Kaiser et al., 2018; Kromdijk et al., 2016). The slow, integrating response of cytosolic NADH/NAD^+^, which does not follow individual fluctuations, suggests that stromal pH changes fast with each oscillation while the cytosolic NAD redox state reflects a longer history. Because the oscillation period also varied during the experiment, the contribution of light history and of the period itself cannot be separated here.

The wavelength dependence of the photosynthetic parameters was systematic and changed with time (Fig. 4B,C and Fig. S3A). The spectrum of variable fluorescence is known to shift during the fast and slow phases of the Kautsky effect, which has been attributed to different chloroplast layers of the leaf contributing at different rates and to the quenching of short-wavelength-emitting chlorophyll by LHCII-associated NPQ (Franck et al., 2005). Heterogeneity between leaf layers and slow regulatory adjustments to the light history (Kaiser et al., 2018) could therefore contribute to the time-dependent differences between detection wavelengths.

Taken together, our methodology opens a new avenue to studying how metabolism and photosynthetic performance are coupled in intact plants. Combined with mutants of the proton circuit or of carbon metabolism, with additional biosensors for other metabolites, or with plants exposed to stress, the approach could help determine which metabolic changes drive, and which follow, the adjustment of photosynthesis to a changing environment. Used together with confocal imaging, as well as other fluorometric approaches and genetic control of sensor expression and subcellular targeting, it can link such whole-plant responses to specific tissues and cells, bridging scales from the cell to the intact plant.

## Material and methods

### Plant material and growth conditions

*Arabidopsis thaliana* (L.) Heynh. (accession Col-0) seeds were stratified for 48 h at 4 °C in the dark and sown in pots with Einheitserde Profisubstrat (Einheitserde Werkverband e.V., Germany). The pots were then transferred to the light and covered with transparent domes for the first 5 days to maintain humidity during acclimation. Plants were watered with tap water every two to three days, grown under long-day conditions (16 h light at 100–120 µE, 22 °C; 8 h dark, 18 °C) and used at 4 to 5 weeks of age. We used stable lines expressing pHluorin2 in the chloroplast stroma, (Cislaghi et al., 2026) ATeam1.03-nD/nA in the stroma (De Col et al., 2017) and Peredox-mCherry in the cytosol (Steinbeck et al., 2020). The line expressing pHluorin2 in the cytosol was generated for this study by transforming Col-0 plants with a construct carrying pHluorin2 under the control of the *UBIQUITIN 10* promoter using the floral dip method, and transformants were selected with hygromycin. All experiments were performed on fully expanded leaves of plants before flowering, which were dark-adapted for 1 h before every measurement on both platforms.

### Confocal laser-scanning microscopy

Leaf discs were mounted between two coverslips and imaged on a Zeiss LSM 980 microscope with a Plan-Apochromat 10×/0.45 M27 objective, focusing on the adaxial mesophyll layer. The microscope was connected to an on-stage illumination device consisting of a ring of LEDs, which exposed the samples to the programmed light regime (Zheng et al., 2025). pHluorin2 was excited at 405 and 488 nm, and emission was collected at 508–535 nm with a GaAsP-PMT detector. The pinhole was set to 120 µm for all channels. Images were acquired every 15 s under red actinic light of 100 µE, with the dark, light and dark phases each lasting 10 min. For Fig. S1, the actinic light was set to 200 and 850 µE with the same phases of 10 min. Time series were processed with the Redox Ratio Analysis (RRA) software (Fricker, 2016), and the pHluorin2 ratio was calculated from the two excitation channels.

### ChloroSpec measurements

Measurements were carried out with a ChloroSpec L1 equipped with the biosensor module (Nanda et al., 2024; Hipsch et al., 2025). A leaf of an intact plant was fixed between two clips of the instrument with the adaxial side up. Emission was detected across 300–900 nm. The actinic LED of the instrument emits at 620 nm, and the other two LEDs were adapted to the sensor. For pHluorin2, the sample was excited with LEDs at 405 and 465 nm in addition to the 620 nm actinic LED, and the ratio was calculated from the fluorescence intensities at 511 nm emission obtained with each excitation. Peredox-mCherry was measured with the 405 LED. The signal was the fluorescence intensity of tSapphire alone, with excitation at 405 nm and emission at 520 nm, and the mCherry channel was not used, so that the response is intensity-based and not ratiometric. For ATeam1.03-nD/nA, the 465 nm LED was replaced by a 440 nm LED, and the sample was excited at 440 nm, with emission measured at 480 and 530 nm.

Illumination protocols were defined in advance and applied automatically. In the dark–light– dark regime of Fig. 1, the actinic light was set to 100 µE, and the dark, light and dark phases each lasted 10 min, as in the CLSM experiments. The same regime was used for cytosolic pHluorin2 and Peredox-mCherry in Fig. 2A,C, whereas for ATeam1.03-nD/nA in Fig. 2B the regime consisted of 5 min of dark, 10 min of light at 100 µE and about 13 min of dark. For the light-intensity series in Fig. 3, leaves were exposed to alternating 5-min dark and light phases of increasing intensity, from 25 to 800 µE (25, 100, 200, 400, 600 and 800 µE). In the fluctuating-light experiment (Fig. 4), recordings started with 10 min of darkness, and data are shown from the onset of illumination: leaves were first illuminated for 15 min at a constant 150 µE and then exposed for about 3.8 h to a sinusoidal modulation between 50 and 250 µE around a mean of 150 µE. The period of the modulation lengthened from about 2 min to about 11 min towards the middle of the series and shortened again to about 3 min, and the experiment ended with a dark phase of about 15 min, during which the interval between measurements increased progressively. Plants expressing Peredox-mCherry were exposed to the same regime for Fig. S3B.

### Photosynthetic parameters

Photosynthetic parameters were calculated from chlorophyll fluorescence recorded at the detection wavelengths 685 nm, 700 nm and 720 nm. At each measuring cycle, the instrument applied a burst of 150 single-turnover flashes (50 + 100 pulses, each 130 µs wide, of 60,000 µE red light, separated by dark intervals of 4 µs) followed by a multi-turnover flash of 15,000 µE and recorded the resulting fluorescence induction curves. The first value of each curve was taken as F_s_, the steady-state fluorescence immediately before the burst, and the last value as the maximum fluorescence (F_m_ in the dark and F_m_′ in the light). F_m_ was taken from the first saturating burst on the dark-adapted leaf. The effective quantum yield of photosystem II, Y(II) = (F_m_′ − F_s_)/F_m_′, and the yields of regulated, Y(NPQ) = F_s_/F_m_′ − F_s_/F_m_, and non-regulated, Y(NO) = F_s_/F_m_, energy dissipation were calculated as described (Genty et al., 1989; Kramer et al., 2004; Klughammer & Schreiber, 2008), and non-photochemical quenching as NPQ = (F_m_ − F_m_′)/F_m_′ (Kramer et al., 2004).

### Data analysis and statistics

Ratios were calculated for each time point and normalized by dividing each value by the mean of the ratio values of the five data points acquired before the light was switched on, and the log_10_ of the normalized ratio is shown. For Peredox-mCherry, the tSapphire intensity was normalized in the same way. Unless stated otherwise, *n* represents independent plants or leaf discs from independent plants. The fluctuating-light experiment (Fig. 4) shows a representative trace from three independent repetitions.

## Declaration of generative AI and AI-assisted technologies in the manuscript preparation process

During the preparation of this work, the authors used ChatGPT (OpenAI) to generate the cartoons of pHluorin2, the CLSM, the light illumination device, ChloroSpec, the data collection system and Arabidopsis plants, and Claude (Anthropic) to edit the language of the text. The authors reviewed and revised all AI-generated content and take full responsibility for the publication.

## Author contributions

Conceptualization: APC, PB, JON, MSchw

Methodology: ARH, APC, MH, AB

Investigation: APC, PB

Visualization: APC

Funding acquisition: MSchw, KBB

Project administration: MSchw, KBB, ARH

Supervision: MSchw, KB, MHip, FB

Writing – original draft: APC

Writing – review & editing: APC, PB, JON, FB, KBB, ARH, MSchw

## Funding

MSchw, KBB, FB and MHip were funded by the Deutsche Forschungsgemeinschaft (DFG) through the Research Unit FOR 5573 “Dynamic Regulation of the Proton Motive Force in Photosynthesis” (project no. 507704013). Confocal microscopy was enabled by the DFG through the infrastructure grant INST211/903-1 FUGG (project no. 429542710).

## Acknowledgements

We thank Jens Appel (University of Kassel, Germany) for providing the pHluorin2 sequence.

## Conflict of interest statement

ARH is a co-founder and Chief Technology Officer (CTO) of ChloroSpec B.V., which manufactures and markets the ChloroSpec L1 spectrometer used in this study. The remaining authors declare no competing interests.

## Notes

### Competing Interest Statement

The authors have declared no competing interest.

### Summary of Updates

Author list and affiliations updated. A new reference was added to the literature. An AI use statement was added.

