## Supplementary Figures 1-3 for "Augmenting chlorophyll fluorescence-based analysis of plant photosynthesis with genetically encoded biosensors"

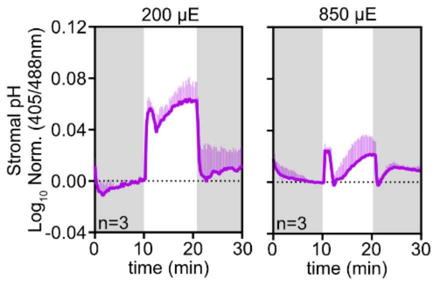


**Figure S1 | Light intensity dependence of stromal pH kinetics measured by CLSM.** Stromal pH measured with pHluorin2 by CLSM in the mesophyll of leaf discs at 200 and 850 µE, as the normalized $\log_{10}$ of the 405/488 nm ratio. Leaf discs were exposed to 10 min of darkness, 10 min of light and 10 min of darkness, with grey shading indicating darkness and white indicating light. Data are mean ± s.d. of *n* = 3 leaf discs, each from a different plant.

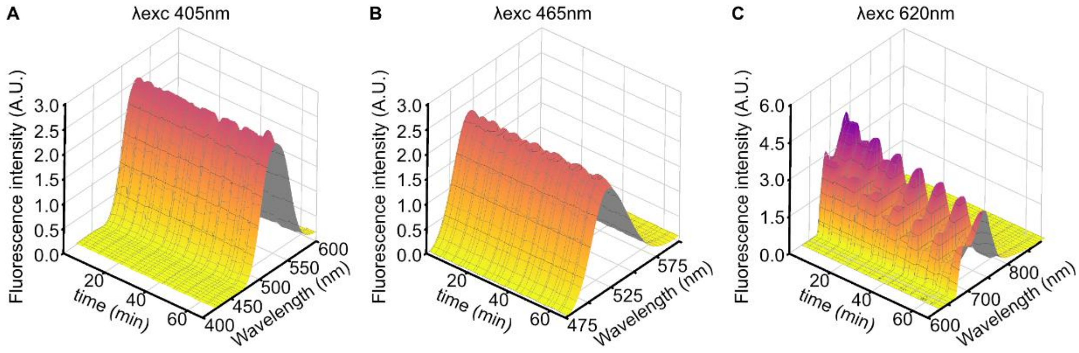


**Figure S2 | Wavelength-resolved fluorescence recorded by ChloroSpec under different excitation LEDs. A,** Emission spectra over time of an attached leaf expressing stromal pHluorin2, recorded between 420 and 600 nm with excitation at 405 nm for biosensor detection, during the light-intensity series shown in Fig. 3. **B,** Emission spectra recorded between 475 and 575 nm with excitation at 465 nm for biosensor detection. **C**, Emission spectra recorded between 620 and 900 nm with excitation at 620 nm for chlorophyll fluorescence detection. Surfaces show fluorescence intensity as a function of time and emission wavelength from one representative leaf.

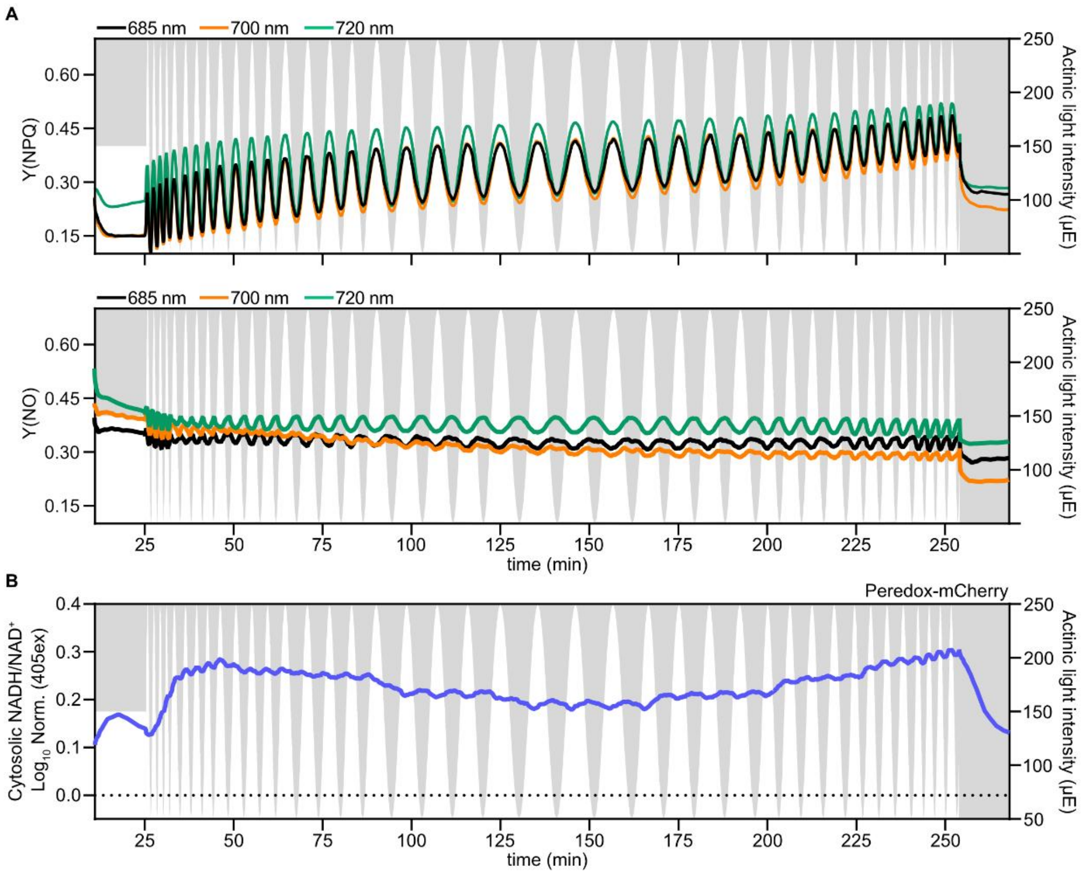


**Figure S3 | Additional photosynthetic parameters and cytosolic NADH/NAD$^+$ under fluctuating illumination. A,** Yield of regulated energy dissipation, Y(NPQ), top, and yield of non-regulated energy dissipation, Y(NO), bottom, calculated from fluorescence at 685 nm, shown in black, at 700 nm, in orange, and at 720 nm, in green, in the same leaf as in Fig. 4. The actinic light intensity is read on the right y-axes, with white regions indicating illumination and grey regions the absence of light. **B,** Cytosolic NADH/NAD$^+$ measured with Peredox-mCherry as the normalized $\log_{10}$ of tSapphire fluorescence, with 405 nm excitation and 520 nm emission, in a separate plant exposed to the light regime in Fig. 4. The trace shows one representative leaf of three independent experiments. Time is counted from the start of the recording, as in Fig. 4.
